# YebC2 resolves ribosome stalling at polyprolines independent of EF-P and the ABCF ATPase YfmR

**DOI:** 10.1101/2024.10.18.618948

**Authors:** Hye-Rim Hong, Cassidy R. Prince, Letian Wu, Isabella N. Lin, Heather A. Feaga

**Affiliations:** Department of Microbiology, Cornell University, Ithaca, NY 14853

## Abstract

Polyproline motifs are essential structural features of many proteins, and recent evidence suggests that EF-P is one of several factors that facilitate their translation. For example, YfmR was recently identified as a protein that prevents ribosome stalling at proline-containing sequences in the absence of EF-P. Here, we show that the YebC-family protein YebC2 (formerly YeeI) functions as a translation factor in *B. subtilis* that resolves ribosome stalling at polyprolines. We demonstrate that YebC2, EF-P and YfmR act independently to support cellular fitness. Moreover, we show that YebC2 interacts directly with the 70S ribosome, supporting a direct role for YebC2 in translation. Finally, we assess the evolutionary relationship between YebC2 and other characterized YebC family proteins, and present evidence that transcription and translation factors within the YebC family have evolved separately. Altogether our work identifies YebC2 as a translation factor that resolves ribosome stalling and provides crucial insight into the relationship between YebC2, EF-P, and YfmR, three factors that prevent ribosome stalling at prolines.

## Introduction

Ribosomes catalyze peptide bond formation between amino acids to produce proteins. The polymerization rate is heavily influenced by the identity of the amino acids involved, with proline posing a special challenge due to its side chain forming a rigid pyrrolidine loop that limits flexibility of the peptide backbone in the ribosomal exit tunnel [1,2]. EF-P was the first protein shown to resolve ribosome stalling at polyprolines and other difficult-to-translation sequences [1,3–9]. EF-P interacts transiently with the ribosomal E-site and then binds stably when tRNA^Pro^ is present in the P-site [10,11]. EF-P binding promotes a favorable geometry of the polypeptide in the exit tunnel to facilitate peptide bond formation [1,2]. *efp* is essential in *Mycobacterium tuberculosis*, *Acinetobacter baumannii*, and *Neisseria meningitidis* [12–14]. In contrast, *efp* deletion from *B. subtilis* causes sporulation and motility defects but does not cause a growth defect in standard lab conditions [15–19].

Recently, our group and Takada and colleagues identified YfmR as a protein that prevents ribosome stalling at polyproline tracts and Asp-Pro motifs in *Bacillus subtilis,* and which becomes more essential in the absence of EF-P [20,21]. YfmR is a member of the ABCF family of ATPases that are widespread throughout bacteria and eukaryotes and have diverse roles in preventing ribosome stalling and mediating antibiotic resistance [22–27]. The *Escherichia coli* homolog of YfmR, Uup, resolves ribosome stalling at polyprolines *in vitro* [28]. A recent structure of Uup bound to *E. coli* ribosomes reveals that it binds the ribosomal E-site and makes contacts with the peptidyl-transferase center [29,30], suggesting that YfmR/Uup may promote peptide bond formation in a manner similar to EF-P. In support of this model, deletion of *yfmR* or *efp* does not result in a fitness defect in *B. subtilis*, while deletion of *yfmR* and *efp* resulted in a severe synthetic fitness defect [21].

The screen we used to identify YfmR also uncovered *yebC2* (formerly *yeeI*) as a gene that may be important for fitness in Δ*efp* cells. Consistent with this finding, a screen performed by Hummels and colleagues in 2019 also identified *yebC2* (*yeeI*) as a gene whose over-expression could rescue the swarming motility defect of Δ*efp B. subtilis* cells [18]. YebC family proteins are annotated as transcription factors in bacteria since these proteins exhibit promoter binding activity and *yebC* deletion causes differential gene expression in *E. coli*, *Pseudomonas aeruginosa*, *Lactobacillus delbrueckii* and in *Borrelia burgdorferi* [31–34]. The human YebC homolog, TACOI, is localized to mitochondria where it is important for efficient translation of COXI [35,36]. TACO1 was recently shown by mitoribosome profiling to prevent ribosome stalling at XPPX motifs and therefore accelerate translation of COXI in human cells [37] and recent work by Ignatov and colleagues demonstrates that YebC in *Streptococcus pyogenes* (YebC_II) facilitates translation of polyproline motifs both *in vivo* and *in vitro* [38].

Here, we show that *B. subtilis* YebC2 is a translation factor that prevents ribosome stalling at a polyproline tract and determine its genetic interaction with *efp* and *yfmR*. Depleting EF-P from Δ*yebC2* cells causes a severe fitness defect, and this defect is even more severe in Δ*yebC2*Δ*yfmR* cells, suggesting that EF-P, YfmR, and YebC2 function independently to support cellular fitness. We find that cells lacking both EF-P and YebC2 exhibit severe ribosome stalling at a polyproline track *in vivo* and that over-expression of YebC2 in Δ*efp* cells reduces ribosome stalling. We show that YebC2 associates with 70S ribosomes, which suggests that YebC2 facilitates translation by acting directly on the ribosome. Finally, we present evidence that YebC2 proteins represent a class of translation factors that are evolutionarily distinct from the previously characterized YebC transcription factors.

## Results

### Deletion of *efp* and *yebC2* causes a severe growth defect and impaired protein synthesis

Previously, we investigated genetic interactions with *efp* using Tn-seq [20]. This screen predicted that *yfmR* is more essential in the absence of EF-P, and we confirmed this result with CRISPRi [20]. Additional genes that may be more essential in the absence of EF-P included genes encoding proline and glycine tRNAs, which decode codons at which ribosomes are more prone to stall in the absence of EF-P (Table 1). Our Tn-seq screen also identified *yebC2* (*yeeI*) as a gene that may be important for growth in the absence of *efp*. *yebC2* exhibited 4-fold fewer insertions in Δ*efp* compared to wild type (Table 1). To test whether YebC2 is more essential in the Δ*efp* background we deleted *yebC2* from Δ*efp* cells. Growth of this strain is severely impaired compared to Δ*efp* or Δ*yebC2* single deletions (Fig. 1A). We complemented this growth defect by providing a single copy of *yebC2* integrated into the chromosome under the control of an IPTG-inducible promoter (Fig. 1A). Moreover, Δ*efp*Δ*yebC2* cells also exhibit a severe decrease in polysomes consistent with a defect in protein synthesis (Fig. 1B).

**Figure 1.**
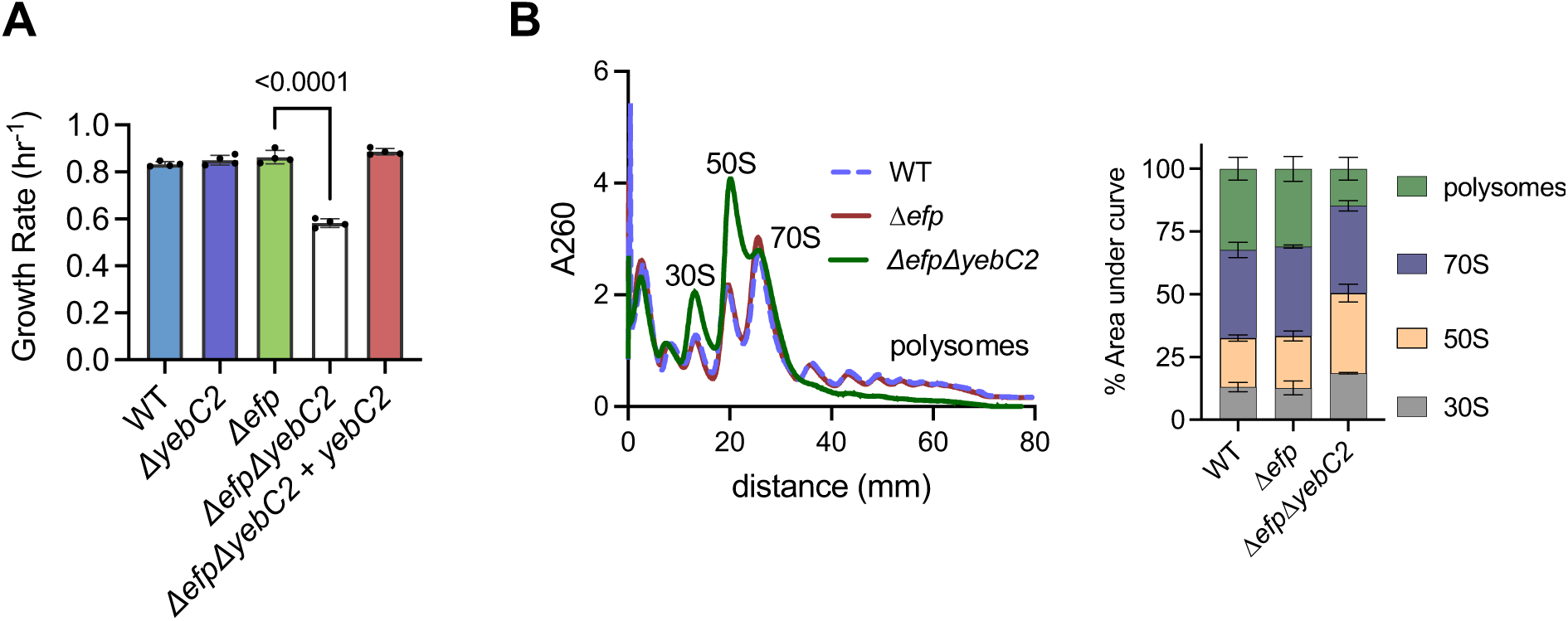
Loss of *efp* and *yebC2* results in severe growth and fitness defects. **(A)**Growth rates of Δ*efp* and Δ*efp*Δ*yebC2* grown in LB at 37°C. The growth defect is complemented by expressing YebC2 from a *hyperspank* promoter (Δ*efp*Δ*yebC2* + *yebC2*). Error bars represent standard deviation of three independent experiments and p-values represent results of an unpaired t-test with Welch’s correction. **(B)** Polysome profiles of wild-type, Δ*efp*, and Δ*efp*Δ*yebC2* strains. Representative of three independent experiments is shown. Quantification shows relative abundance of each ribosomal species as determined by area under each curve. Error bars represent standard deviation of three independent experiments.

**Table 1.** Selected genes identified as interacting with EF-P by Tn-seq. Avg. insertions in each background

| <u>Gene</u> | <u>Wild type</u> | <u>$\Delta efp$</u> | <u>Log<sub>2</sub>(fold change)</u> |
| --- | --- | --- | --- |
| *yfmR* | 147 | 12 | – 3.6 |
| *trnJ-Pro* | 50 | 7 | – 2.8 |
| *trnI-Pro* | 44 | 7 | – 2.6 |
| *yebC2* (*yeel*) | 796 | 192 | – 2.1 |

**Table 2.**
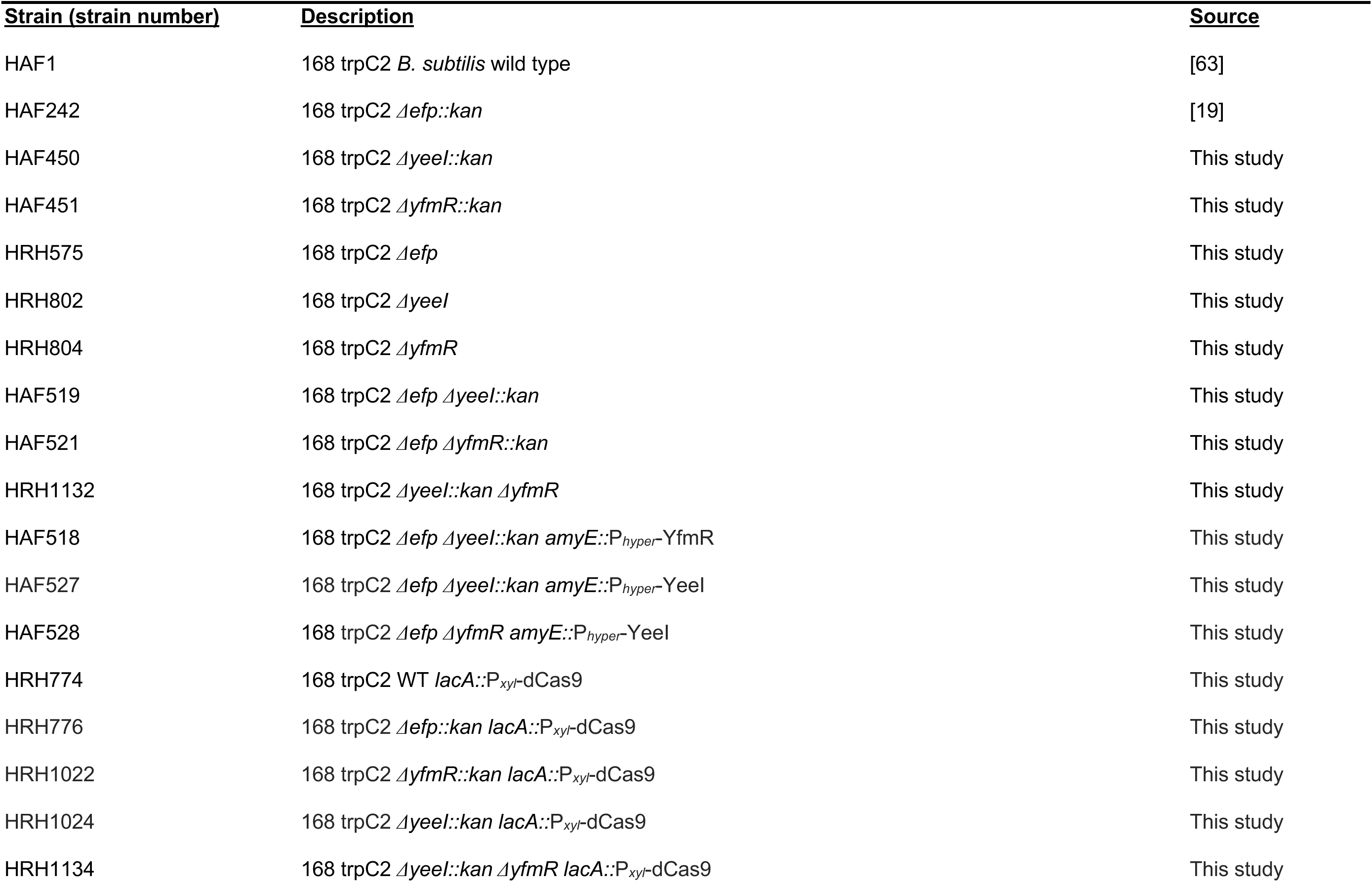
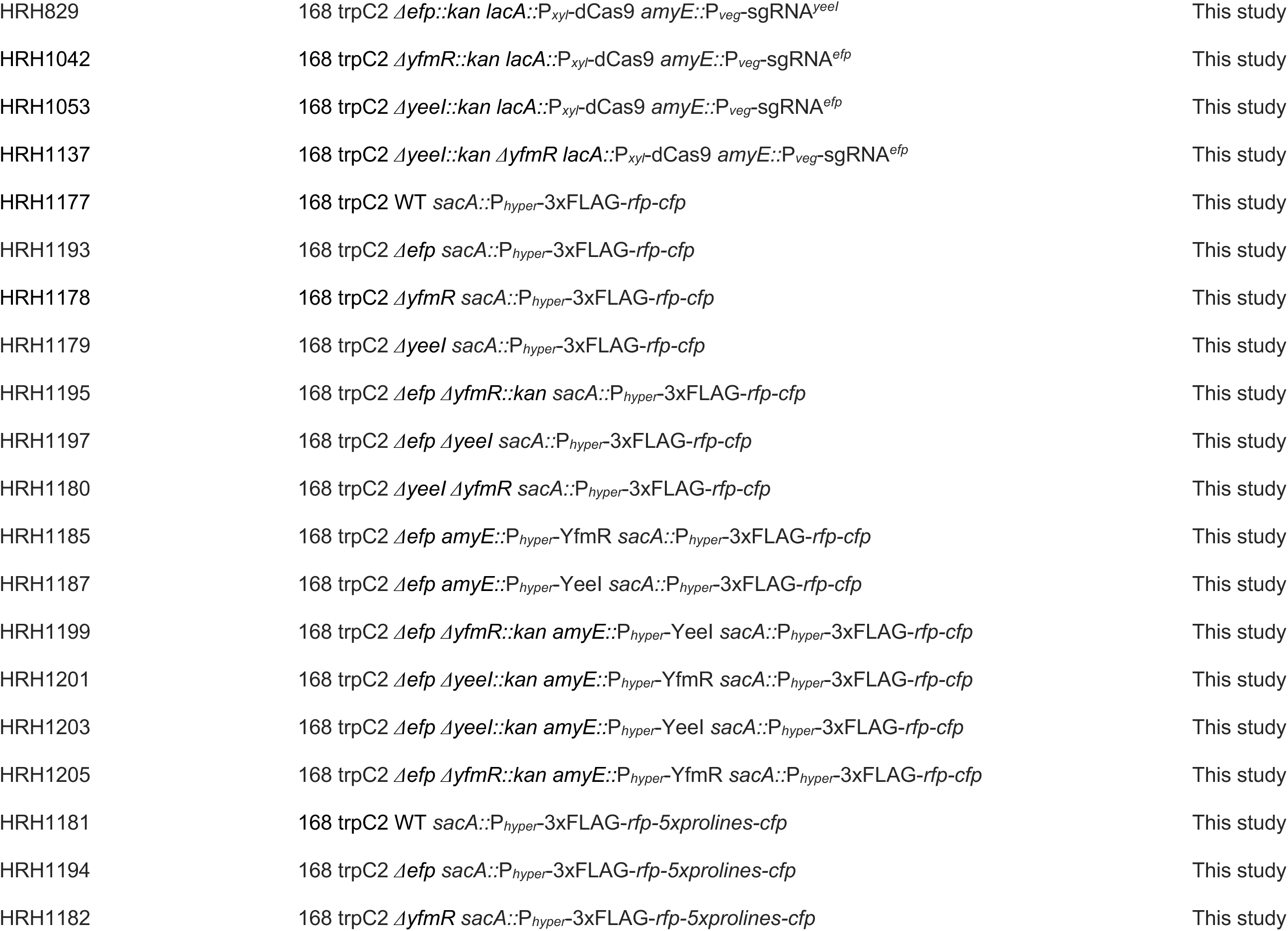
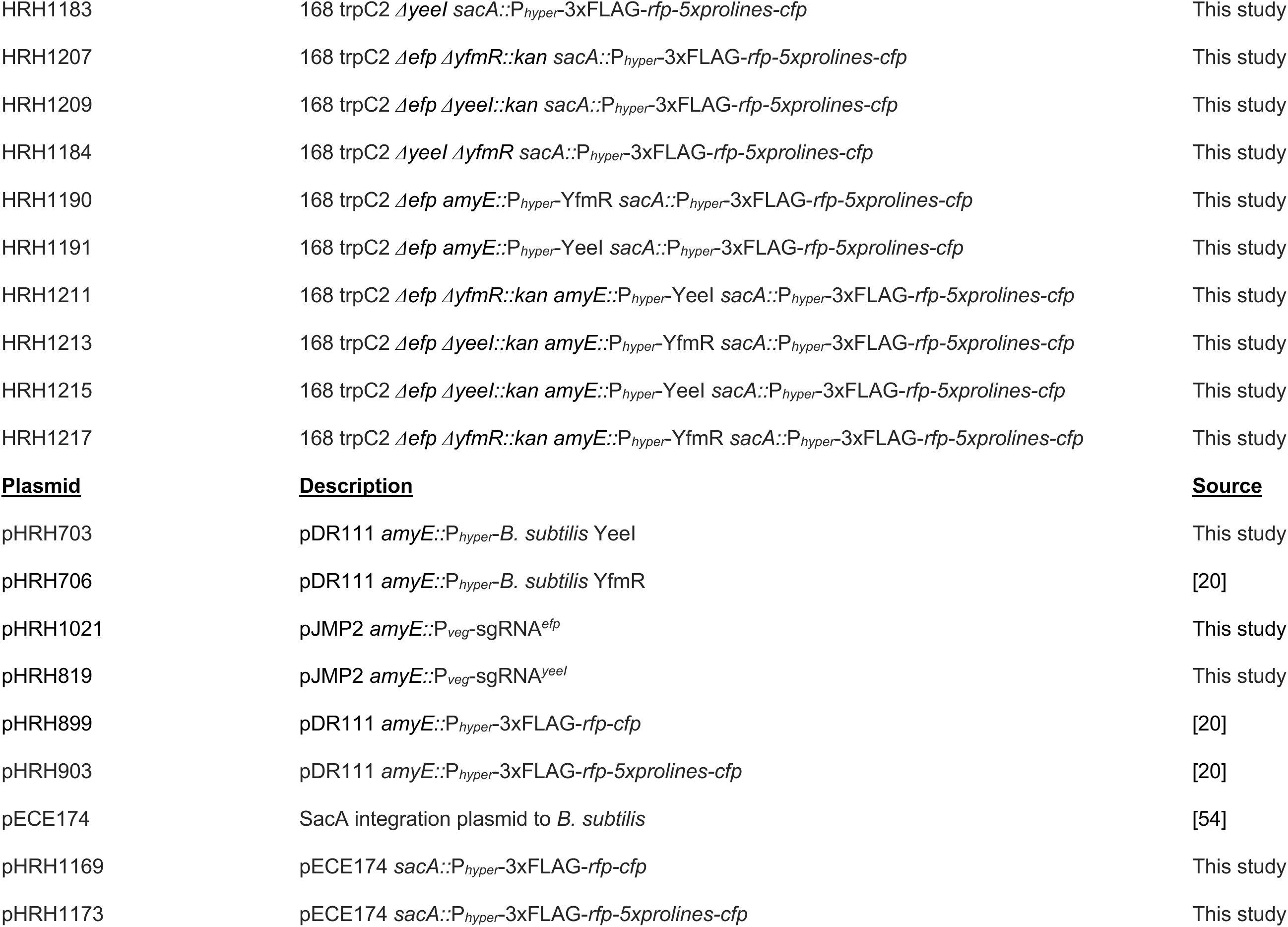
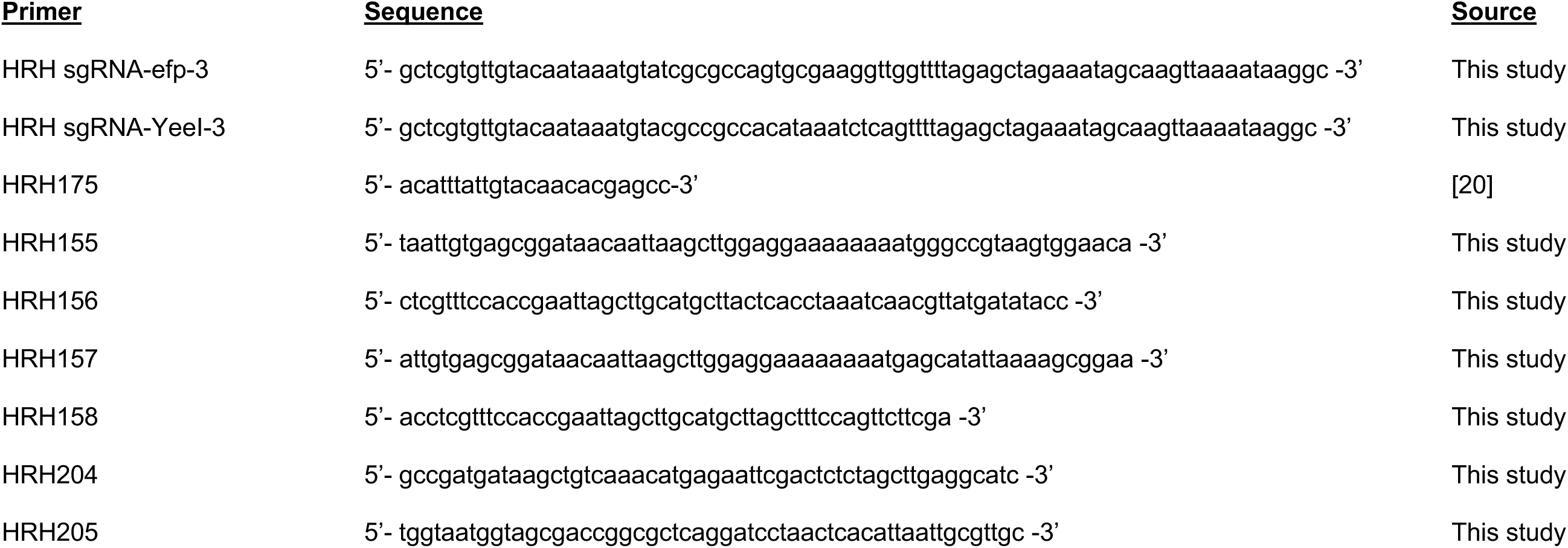

### YebC2 is important for cellular fitness in the absence of EF-P and YfmR

We next tested whether EF-P depletion from Δ*yfmR*Δ*yebC2* cells would cause a more severe fitness defect than depletion of EF-P from Δ*yfmR* or from Δ*yebC2* single deletions using CRISPR interference [39]. We constructed a strain expressing a guide RNA (sgRNA*^efp^*) that blocks transcription of *efp* when expressed alongside a deactivated Cas9 (dCas9) [39,40]. We then prepared dilutions of these cells and plated them with and without xylose to induce dCas9 (Fig. 2A). Consistent with our previous observations, depleting *efp* from Δ*yfmR* cells decreased colony formation by 3 orders of magnitude compared to when dCas9 was not induced. When *efp* was depleted from Δ*yfmR*Δ*yebC2* cells, colony formation decreased by 4 orders of magnitude (Fig. 2A). We did not detect any difference in growth or survival of Δ*yfmR*Δ*yebC2* double deletion. These results suggest that YebC2 is important in the Δ*efp* background, and even more important in a strain lacking both EF-P and YfmR.

**Figure 2.**
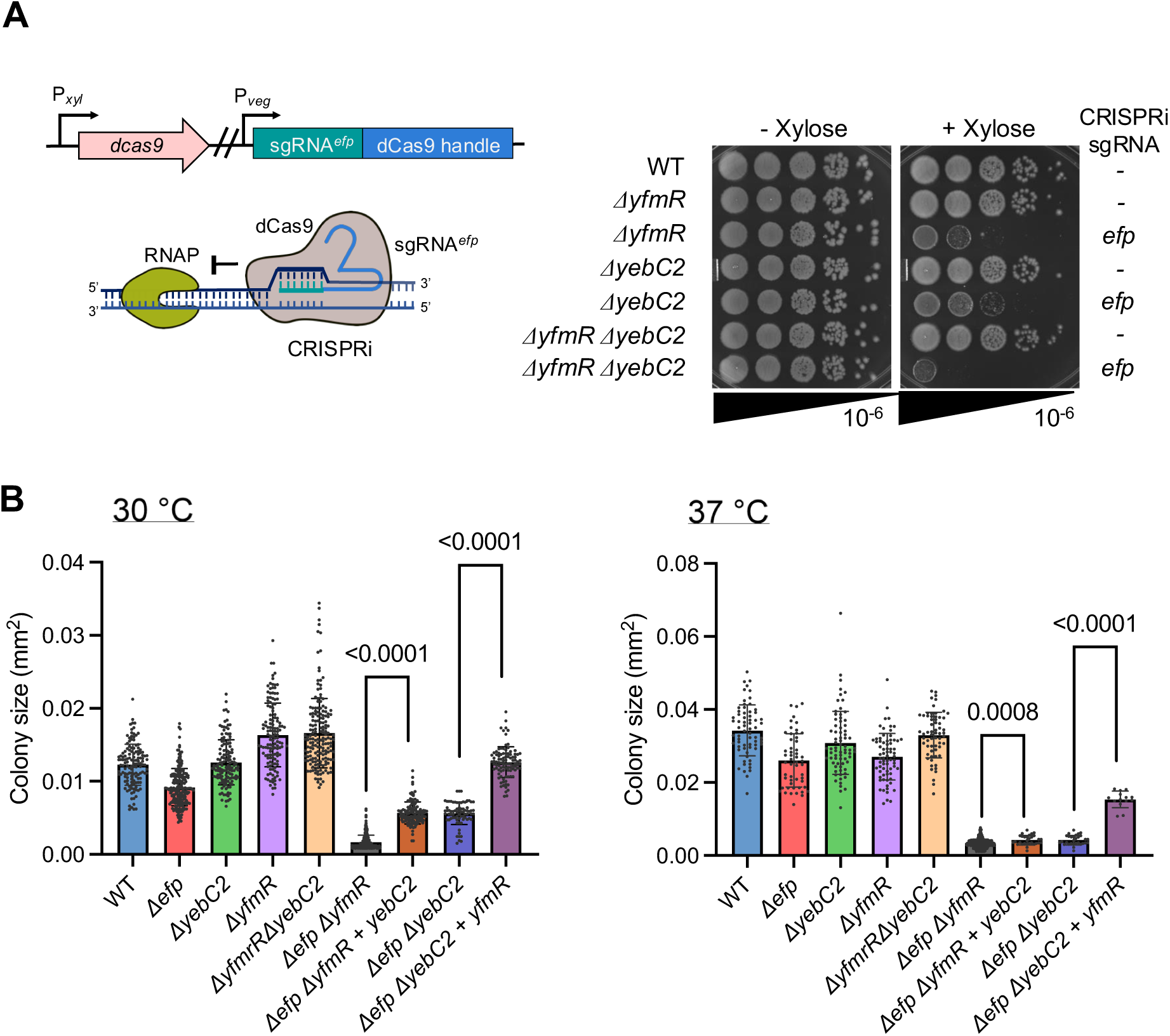
Ectopic expression of YebC2 significantly increases fitness of Δ*efp*Δ*yfmR* cells. **(A)**CRISPR interference was used to deplete EF-P from Δ*yfmR*, Δ*yebC2*, or Δ*yfmR*Δ*yebC2* double deletion. Culture was serially diluted and plated on LB with and without xylose to induce expression of dCas9. **(B)** Colony area measurements indicate that expression of YebC2 in Δ*efp*Δ*yfmR* or expression of YfmR in Δ*efp*Δ*yebC2* cells partially rescues growth. LB plates were incubated at either at 30°C or 37°C. Area of resulting colonies was quantified with ImageJ. Error bars represent standard deviation and p-values represent results of an unpaired t-test with Welch’s correction.

### YebC2 over-expression rescues the synthetic fitness defect of *ΔefpΔyfmR*

Previously, we found that deletion of *yfmR* in *B. subtilis* Δ*efp::mls* is lethal [20]. However, removal of the erythromycin resistance marker allows construction of the Δ*efp*Δ*yfmR* strain, but with a severe a severe synthetic growth defect (Fig. 2B) [21]. Therefore, we tested whether over-expression of YebC2 could rescue this synthetic defect. We expressed YebC2 under the control of an IPTG-inducible promoter in *ΔefpΔyfmR* cells. At both 30°C and 37°C over-expression of YebC2 significantly improves fitness, as determined by colony size measurements (Fig. 2B). We next tested whether YfmR over-expression could rescue growth of Δ*efp*Δ*yebC2* cells. Indeed, expression of YfmR in *ΔefpΔyebC2* cells also rescued growth as determined by colony size measurements (Fig. 2B). Since EF-P depletion from Δ*yfmR*Δ*yebC2* was more severe than depletion from either single mutant, and since over-expression of YebC2 or YfmR in the absence of the other two factors partially rescues growth, we conclude that YebC2, YfmR, and EF-P act independently to support growth.

### YebC2 prevents ribosomal stalling at polyprolines

To determine whether YebC2 is important for preventing ribosome stalling at polyprolines, we used an *in vivo* stalling reporter encoding an N-terminal Flag tag for detection and five consecutive prolines mid-way through the protein sequence (Fig. 3A). If ribosomes stall at the polyproline tract a truncated peptide is produced. Percent stalling was quantified as a percentage of stalled peptide divided by the sum of the stalled plus full-length peptide.

**Figure 3.**
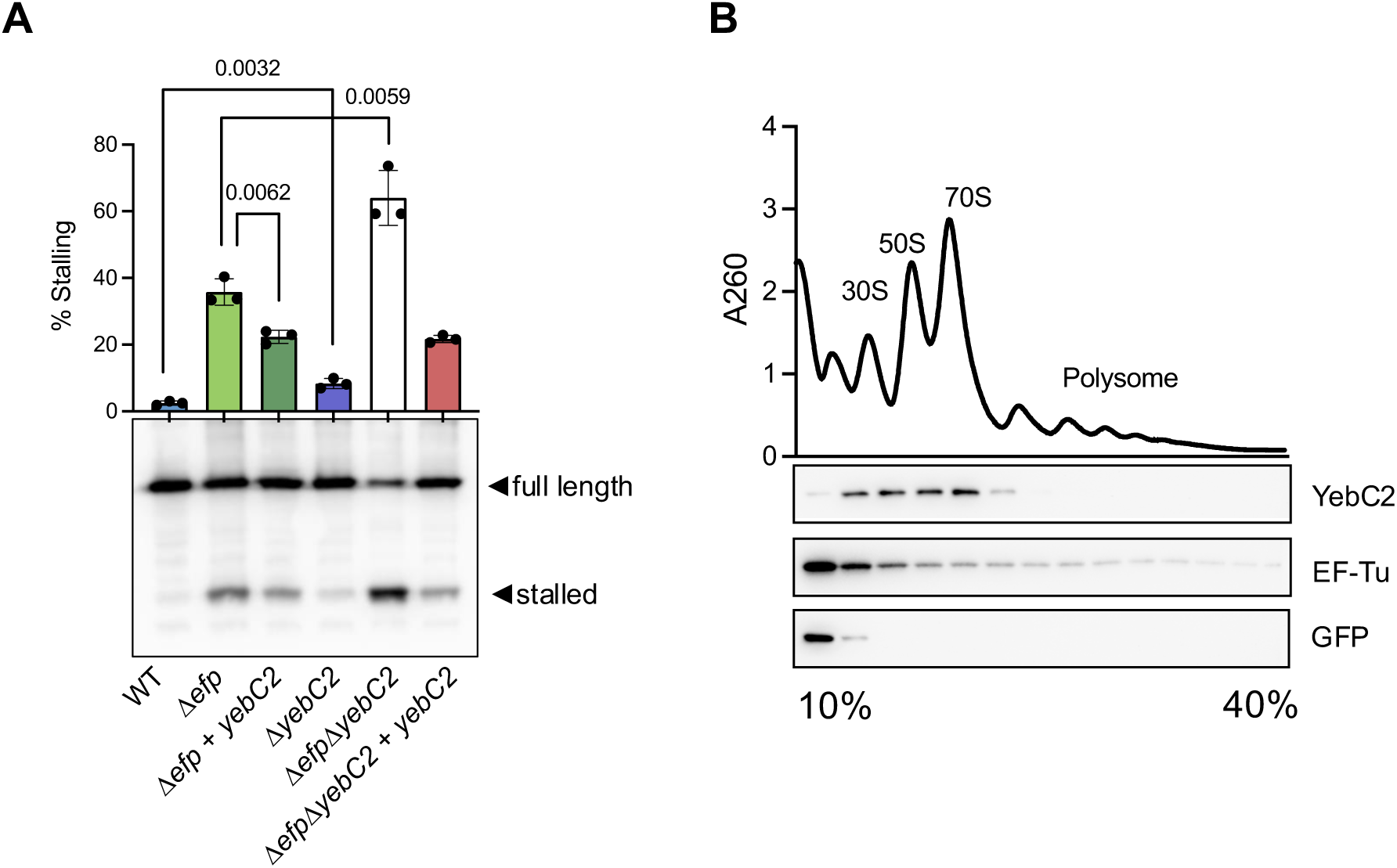
YebC2 prevents ribosome stalling at a polyproline tract *in vivo* and associates with 70S ribosomes. **(A)** A reporter encoding a penta-proline tract was used to monitor ribosome stalling *in vivo*. Percent stalling is reported as level of stalled protein divided by the sum of stalled and full-length protein. P-values are the result of an unpaired t-test performed on three biological replicates. Error bars indicate standard deviation. **(B)** Lysate from a strain expressing His-tagged YebC2 was resolved by sucrose density gradient ultracentrifugation. Fractions were probed with anti-His antibody or a polyclonal antibody raised against EF-Tu. His-tagged GFP was used as a negative control for ribosome association.

*ΔyebC2* cells exhibit ribosome stalling that is significantly higher than in wild-type cells (p = 0.0032)(Fig. 3A). The ribosome stalling observed in Δ*yebC2* cells (8 ± 1%) is not as high as in Δ*efp* cells (36 ± 4%). However, *ΔefpΔyebC2* exhibit very high levels of ribosome stalling (64 ±8%), significantly higher than cells deleted for just *efp* (p = 0.0059). Moreover, YebC2 over-expression in *Δefp* cells significantly reduced ribosome stalling (p = 0.0062). The additive increase in ribosome stalling for the *ΔefpΔyebC2* double deletion and decreased ribosome stalling when YebC2 is over-expressed in the absence of EF-P further demonstrates that these proteins can function independently to prevent ribosome stalling. Altogether, these results demonstrate that YebC2 prevents ribosome stalling at polyproline tracts.

### YebC2 associates with 70S ribosomes

To determine whether YebC2 directly acts on the ribosome, we constructed a His-tagged version of YebC2 to monitor ribosome association. His-tagged YebC2 was functional, as evidenced by its ability to complement the impaired growth of Δ*efp*Δ*yebC2* cells (Fig S1). Cells expressing His-tagged YebC2 were harvested in late exponential phase, lysed, and cleared lysate was resolved on by sucrose density gradient ultracentrifugation. We found that YebC2 co-migrates with ribosomes and was strongly associated with 70S ribosomes (Fig. 3B). In contrast, a His-tagged GFP that served as a negative control for ribosome association was found only at the top of the gradient. These results suggest that YebC2 exerts its anti-stalling activity by acting directly on the ribosome.

### YebC2 is evolutionarily distinct from YebC transcription factors

Many bacterial species encode two YebC paralogs [41]. Since most YebC paralogs studied to date are characterized transcription factors, we asked whether YebC2 clades separately from these factors. To determine the evolutionary relationship between the YebC paralogs we built a maximum likelihood tree based on the protein sequences of >15,000 YebC family proteins (Fig. 4). We found that the YebC paralogs that have experimental support for a role in transcription (YebC from *E. coli*, *L. delbrueckii, B. burgdorferi* and PmpR from *P. aeruginosa*) cluster together, while those that have a role in translation (*B. subtilis* and *S. pyogenes* YebC2 and *E. coli* YeeN) cluster separately (99.9% maximum likelihood bootstrap value) (Fig. 4). YebC2 from *B. subtilis* and *S. pyogenes* share a common ancestor exclusive of the YebC proteins that have been characterized as transcription factors (100% maximum likelihood bootstrap value) (Fig. S2). Importantly, this clustering is not based on species phylogeny, since *B. subtilis* YrbC clusters with the transcription factors. Consistent with this clustering, our Tn-seq screen did not identify a genetic interaction between *yrbC* and *efp* [20]. These data suggest that YebC-family proteins have evolved separately to function in transcription or translation.

**Figure 4.**
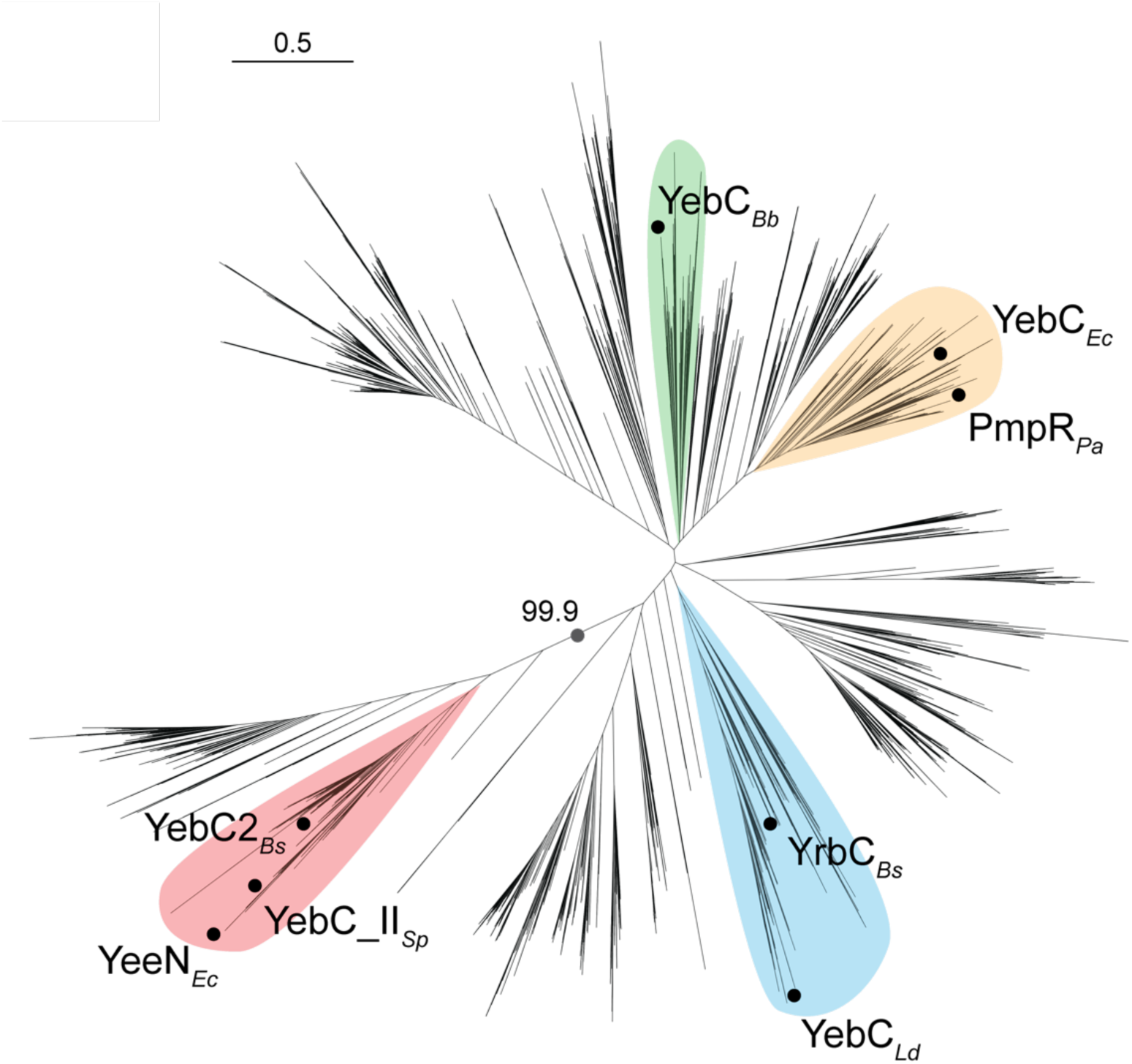
YebC transcription factors are evolutionarily distinct from YebC2 translation factors. The unrooted maximum-likelihood tree was built using all YebC family protein sequences detected in a database of >15,000 prokaryotic representative genomes. Characterized YebC family proteins are denoted with a circle and labeled with their given gene name and respective organism: *Bs, Bacillus subtilis; Ld Lactobacillus delbrueckii; Pa, Pseudomonas aeruginosa; Ec, Escherichia coli; Bb, Borrelia burgdorferi; Sp, Streptococcus pyogenes.* Clades containing characterized proteins were highlighted.

### YebC proteins are widely distributed in bacteria while YebC2 proteins are more restricted

Having determined that YebC and YebC2 proteins are evolutionarily distinct, we next determined the conservation of these proteins across the bacterial domain (Fig. 5). 87% of the >15,000 bacterial genomes we surveyed encode at least one YebC-family protein, consistent with a previous report of the conservation of this protein family [41]. However, YebC is much more widely distributed and highly conserved than YebC2. We detected YebC in 80% of our surveyed genomes and YebC2 in only 13%. YebC2 was mainly restricted to Firmicutes (Bacillota) and Gamma-proteobacteria. Interestingly, we found that some genomes encode up to 3 YebC paralogs and up to 2 YebC2 paralogs (Fig. 5). Further work is needed to determine whether YebC2 it is advantageous to have more than one copy of YebC2 or whether each of the YebC2 proteins has any specialized function in rescuing stalled ribosomes.

**Figure 5.**
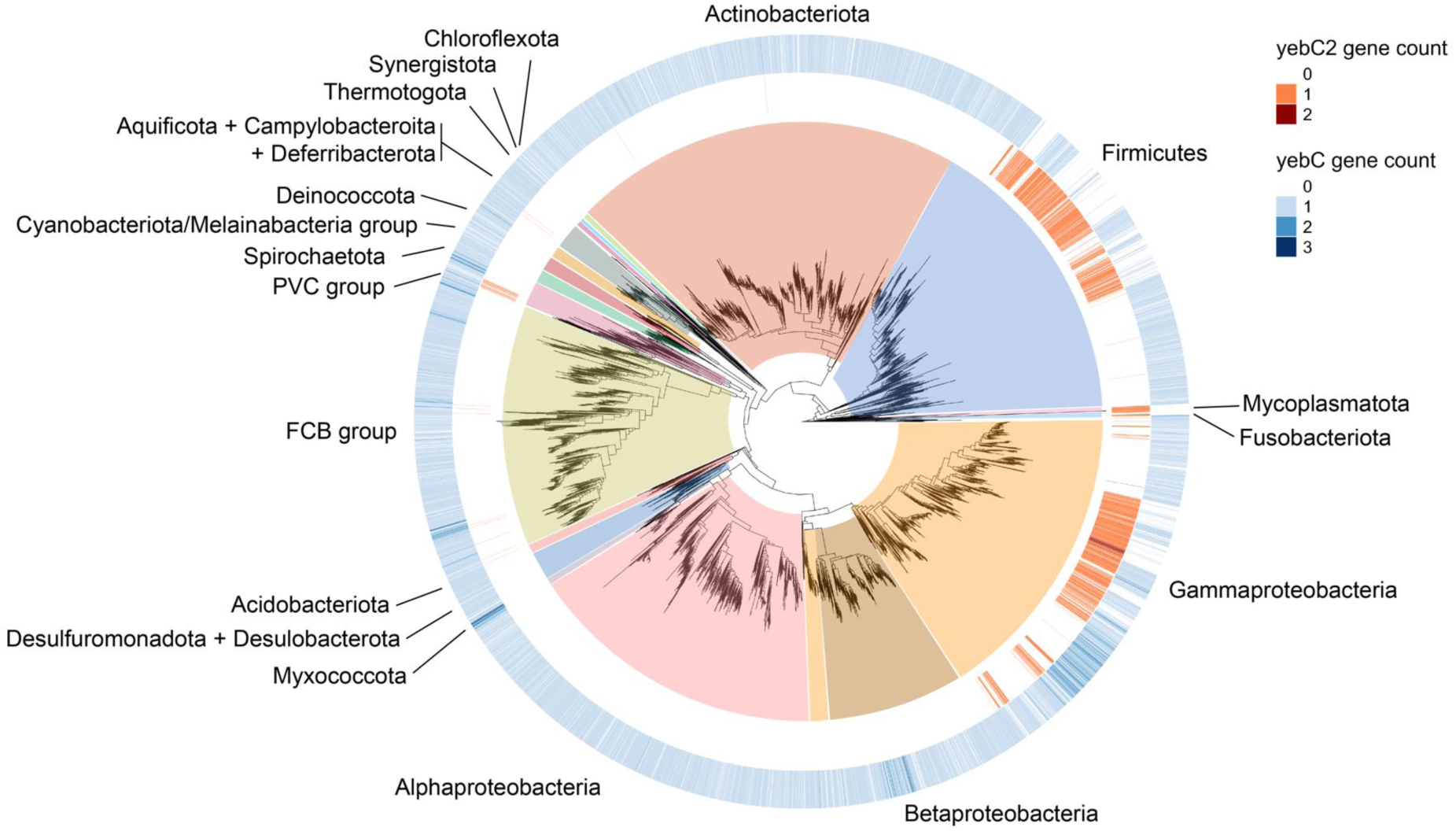
Distribution of YebC-family paralogs in bacteria. A midpoint rooted 16S maximum-likelihood phylogenetic tree of species in the bacterial domain, indicating the number of YebC2 or YebC paralogs in each genome. YebC2 paralogs are most well-conserved in Firmicutes (Bacillota), and Gamma-proteobacteria whereas YebC paralogs are widely distributed across most bacterial phyla. 87% of surveyed genomes encoded at least on YebC-family protein.

## Discussion

Here we show that YebC2 is a ribosome-binding protein that resolves ribosome stalling at a penta-proline tract *in vivo* (Fig. 3). These findings are in complete agreement with the elegant work of Brischigliaro and colleagues and Ignatov and colleagues which have likewise determined a role for human TACO1 and *S. pyogenes* YebC_II in preventing ribosome stalling at polyprolines [37,38]. Moreover, we show that simultaneous loss of YebC2, EF-P, and YfmR severely reduces the viability of *B. subtilis,* and present evidence that YebC2 prevents ribosome stalling independent of EF-P and YfmR (Fig. 2). These data contribute to a more complete understanding of the various factors that prevent ribosome stalling at polyproline tracts.

YebC family proteins have been widely annotated as transcription factors [31–34,42,43]. By analyzing YebC and YebC2 protein sequences, we found that these proteins cluster into divergent clades in good agreement with experimental evidence supporting their roles in either transcription or translation (Fig. 4). In particular, *B. subtilis* and *S. pyogenes* YebC2 share a common ancestor that is exclusive of the YebC transcription factors. However, it is notable that *E. coli* YebC resolves ribosome stalling at a penta-proline tract despite its role in transcription [34] and the fact that it clusters with the YebC transcription factors (Fig. 4). Do YebC transcription factors also play a role in translation? While it is clear that YebC and YebC2 are evolutionarily distinct, more work is needed to fully characterize their functional divergence.

We also determined the conservation of YebC and YebC2 paralogs across the bacterial domain. YebC2 is conserved primarily within Firmicutes (Bacillota) and Gamma-proteobacteria (Fig. 5). YebC is more broadly distributed, with homologs detected in most phyla. The retention of both YebC and YebC2 paralogs in many taxa further supports a model in which these proteins impart unique selective advantages due to independent functions. It is notable that although the transcription-family YebC proteins are broadly distributed in bacteria, eukaryotes encode only the YebC2-type protein [41].

Our data support a model in which EF-P, YfmR, and YebC2 can each function independently to prevent ribosome stalling. An independent role for YebC2 is demonstrated by its ability to prevent stalling on an *in vivo* reporter in Δ*yfmR*Δ*efp* cells and its ability to partially complement the severe synthetic growth defect of Δ*yfmR*Δ*efp* cells (Fig. 2). If YebC2 were absolutely dependent on either EF-P or YfmR for its activity, it would be unable to rescue growth or prevent ribosome stalling in the Δ*efp*Δ*yfmR* background. Interestingly, while EF-P and YfmR have partially overlapping binding sites in the ribosomal E-site, some evidence suggests that YebC2 may bind the A-site. First, proximity labeling with a TACOI-BirA fusion protein resulted in biotin labeling of A-site adjacent ribosomal proteins [37]. Second, sequencing of RNA cross-linked to YebC_II in *S. pyogenes* revealed likely contacts with Helix 89 [38]. If YebC2 and its homologs can bind the ribosomal A-site while EF-P or YfmR are bound in the E-site, then it remains an exciting possibility that there are some forms of ribosome stalling where YebC2 does indeed aid the activity of either EF-P or YfmR.

Although we observe anti-stalling activity for EF-P, YfmR, and YebC2 on a penta-proline reporter, it is likely that all of these factors prevent ribosome stalling at sequences that extend beyond prolines. For example, YfmR also prevents ribosome stalling on polyacidic residues [21]. Meanwhile, EF-P promotes peptide bond formation at other difficult-to-translate sequences [44,45]. In particular, EF-P likely plays a role in formation of the first peptide bond since it recognizes both tRNA^Pro^ and initiating tRNA^fMet^ in the P-site [11,46]. Moreover, EF-P promotes peptide bond formation between initiating formyl-methionine and the second amino acid and helps maintain the reading frame during early elongation [47–49]. There is some evidence that YfmR may also participate in early elongation since YfmR depletion in Δ*efp* cells causes increased association of initiator tRNA with stalled ribosomes [20]. There is also evidence that YebC2 plays a role at non-proline encoding sequences since deletion of the YebC2 ortholog in yeast causes a more general defect in protein synthesis, with reduced overall synthesis of mitochondrial-localized reporters [50].

Although structurally distinct, both EF-P and YfmR make similar contacts with the P-site tRNA, and both exhibit tRNA mimicry, which is common for ribosome-binding proteins [51]. The YebC2 predicted structure does not align well with either EF-P or YfmR and so its mechanism of action is likely to be completely different, especially if it binds the ribosomal A-site. Nevertheless, proximity labeling with TACOI and RNA crosslinking with YebC_II indicate that these proteins likely contact the peptidyl-transferase center, and may therefore directly stimulate peptide bond formation [37,38]. Structural studies of YfmR on a polyproline substrate, and of YebR bound the ribosome are essential to our understanding of how these proteins resolve ribosome stalling.

## Materials and Methods

### Strains and media

Strains were derived from *B. subtilis* 168 trpC2 and are listed in Table 1. Single deletions were obtained from the BKK collection [52] and moved into the lab’s 168 trpC2 strain by natural transformation. The kanamycin resistance cassette was excised to make clean deletions using pDR244 [52]. *B. subtilis* strains were cultured in LB and supplemented with antibiotics at final concentrations of 100 µg/mL spectinomycin, 1x MLS (1 µg/mL erythromycin and 25 µg/mL lincomycin), or 5 µg/mL chloramphenicol. *E.coli* DH5alpha strains were cultured in LB with 100 µg/mL ampicillin.

### Complementation of *yebC2* and *yfmR*

Primers are listed in Table 1. *yeeI* was amplified from the wild-type *B. subtilis* 1772 WT 168 trpC2 genomic DNA using primers HRH155 and HRH156 which contain 22 bp of homology to pDR111. Primers HRH157 and HRH158 were used to amplify *yfmR*. The resulting fragments were cloned by Gibson assembly into pDR111 cut with HindIII and SphI. The resulting plasmids, pHRH703 (P*hyper*-*yeeI*) and pHRH706 (P*hyper*-YfmR) were linearized with ScaI and transformed for integration on the chromosome at *amyE*.

### Growth Curves

*B. subtilis* strains were grown overnight at room temperature, and inoculated to a final OD600 0.05 in 150 µl LB, and supplemented with 1 mM IPTG where appropriate in a 96 well-plate (ThermoScientific 167008). The cultures were incubated at 30 °C and 37 °C with linear shaking (2-mm intensity). OD600 of strains were measured at 15-min intervals over 20 hours using a microplate reader (BioTek).

### Colony Size Measurement

*B. subtilis* strains were cultured in LB at room temperature or 37 °C overnight in a roller drum at 80 rpm. 1 mM IPTG was added to the strains overexpressing YeeI or YfmR. The cells were normalized to OD600 0.05 and serially diluted and plated onto two LB agar plates. These plates were incubated at 30 °C or 37 °C for 24 hours, and then placed at room temperature for 24 hours. The plates were imaged on ChemiDoc^TM^MP (Biorad), and the area of the individual colonies was measured using ImageJ [53].

### CRISPRi Depletion

Primer HRH sgRNA-efp-3 containing an sgRNA sequence (5’-tcgcgccagtgcgaaggttg-3’) was designed to target EF-P. HRH sgRNA-efp-3 and HRH175 [20] were used to amplify pJMP2 [40], generating pHRH1021. pJMP1 carrying dCas9 under the xylose-inducible promoter [40] was transformed into the single deletion strains *ΔyeeI::kan* (HAF450) and *ΔyfmR::kan* (HAF451) and the double deletion strain *ΔyeeIΔyfmR* (HRH1132). Next, pHRH1021 was transformed into the strains harboring dCas9, therefore producing EF-P depletion strains. The resulting *B. subtilis* CRISPRi knockdown strains were cultured overnight without xylose and diluted to an OD600 of 0.05 in LB. The cultures were subsequently diluted 10-fold as 10^-2^ to 10^-6^ and spotted onto LB agar without xylose or onto LB agar containing 5% xylose.

### Proline Stalling Reporter and Western Blot

The RFP-CFP fusion cassette containing the pentaproline stalling motifs (5’-ccaccaccaccaccc-3’) or the reporter cassette without the motifs were amplified by using primers HRH204 and HRH205 from the previous constructs pHRH899 and pHRH903 (Table 1). The resulting fragments were cloned into the empty pECE174 [54] plasmid cut with EcoRI and BamHI, producing pHRH1169 and pHRH1173. The resulting reporter plasmids were sequenced by Plasmidsaurus and linearized with ScaI to transform into the different combinations of deletions in *B. subtilis* for recombination at *sacA*. The reporter strains were grown overnight and then diluted back to OD600 0.05. The diluted cultures were induced with 1 mM IPTG and grown up to OD600 1.2 at 37 °C in a roller drum at 90 rpm. 1-mL cell cultures were collected and resuspended with 60 µL of lysis buffer (10 mM Tris pH 8, 50 mM EDTA, 1 mg/mL lysozyme), then incubated at 37 °C for 10 min. Next, the cell lysates were resuspended with 40 µL 4x SDS-PAGE loading buffer. The lysate samples were heated at 85 °C for 5 min and immediately cooled on ice. 10 µL samples were loaded onto a 12 % SDS-PAGE gel and run at 150 V for 70 min. The protein was transferred to PVDF membrane (Biorad) at 300 mAmp for 100 min. The membrane was blocked with 3 % BSA for 20 min and incubated with 4 µg anti-FLAG monoclonal antibody (Sigma SAB4200119) for 20 min at room temperature. The membrane was washed 3 times with PBS-T and developed with ECL (Biorad170-5060) for 2 min and imaged on ChemiDoc^TM^MP (Biorad).

### Polysome Profiling

Strains were grown overnight at 37 °C and inoculated to an OD600 of 0.05 in 40 ml LB the next morning. Cells were collected at OD600 1.2 by centrifugation at 8000 rpm for 10 minutes (Beckman Coulter Avanti J-15R, rotor JA-10.100). Cell pellets were resuspended in 200 µl gradient buffer containing 20 mM Tris (pH 7.4 at 4°C), 0.5 mM EDTA, 60 mM NH4Cl, and 7.5 mM MgCl2 and 6mM 2-mercaptoethanol. Cells lysed using a homogenizer (Beadbug6, Benchmark) by five 20 second pulses at speed 4350 rpm with chilling on ice for 2 min between the cycles and clarified by centrifugation at 21,300 rcf for 20 min (Eppendorf 5425R, rotor FA-24x2). Clarified lysates were normalized to 1500 ng/µl and loaded onto 10 – 40% sucrose gradients in gradient buffer and run for 3 hours at 30,000 rpm at 4°C in an SW-41Ti rotor. Gradients were collected using a Biocomp Gradient Station (BioComp Instruments) with A260 continuous readings (Triax full spectrum flow cell). The area under each peak was quantified using Graphpad Prism.

### Gene detection

Genes were detected in a database of >18,000 representative prokaryotic genomes from NCBI RefSeq using HMMER v3.3 (nhmmer) (hmmer.org) with an E-value cutoff of 0.05 and a query of all characterized *yebC*-family gene sequences. Hits were classified as either *yebC* or *yebC2* depending on the gene query that resulted in a higher sequence bit score, and therefore greater homology. Genomes were filtered for <10% CheckM contamination [55], which left us with 15,259 genomes to survey.

### Phylogenetics

16S rRNA sequences of all genomes were identified and acquired using BLAST v2.13.0, aligned using MAFFT v7.453, and applied to FastTree v2.1.11 [56] to infer a maximum likelihood tree [57]. FastTree produces unrooted phylogenies, so the tree was midpoint rooted using the phangorn v2.11.1 package [58]. Taxonomic classification was assigned to genomes using the NCBI Taxonomy database [59] and taxonkit v0.17.0 [60]. Phyla were named using the conventions in Coleman et al. 2021 [61]. The tree was visualized using ggtree v3.12.0 [62]. For the large YebC-family tree, the gene sequences of HMMER hits were translated using a Python script, and the tree was built and visualized using the same pipeline as described previously.

## Acknowledgements

H-RH, CRP, and HAF were supported by NIH R35GM147049. CRP was supported by a Graduate Research Fellowship from the National Science Foundation. We are grateful to Tory Hendry, Vasili Hauryliuk, Katrina Callan, Kevin England, and Daniel Tetreault for feedback on the manuscript.

---

**Figure S1.**
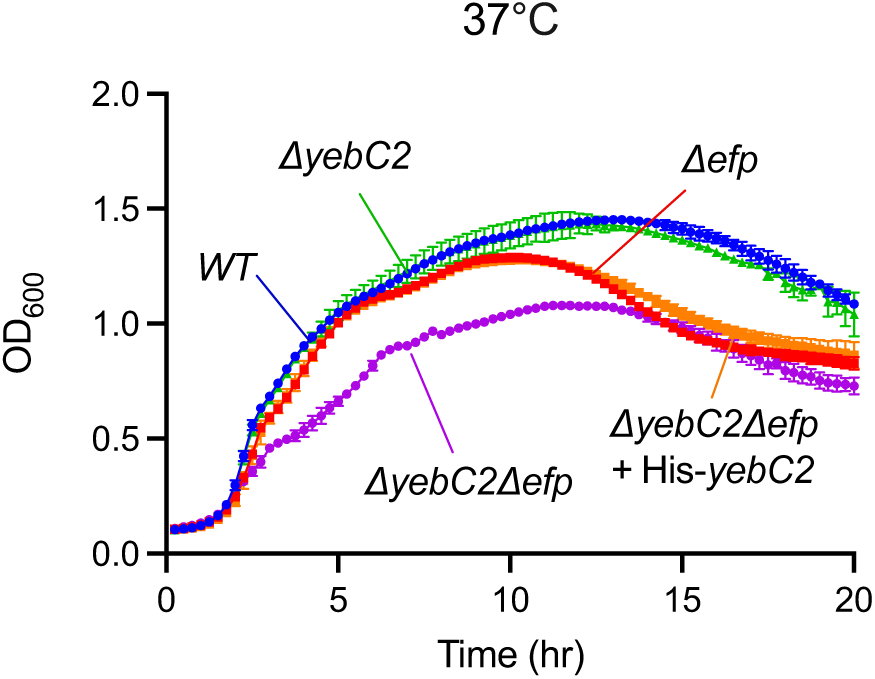
His-Tagged YebC2 is functional and complements the growth defect of Δ*efp*Δ*yebC2* cells in vivo. Growth curves in LB at 37°C are shown for WT, Δ*efp*, Δ*yebC2*, Δ*yebC2*Δ*efp* and Δ*yebC2*Δ*efp* expressing His-tagged YebC2.

**Figure S2.**
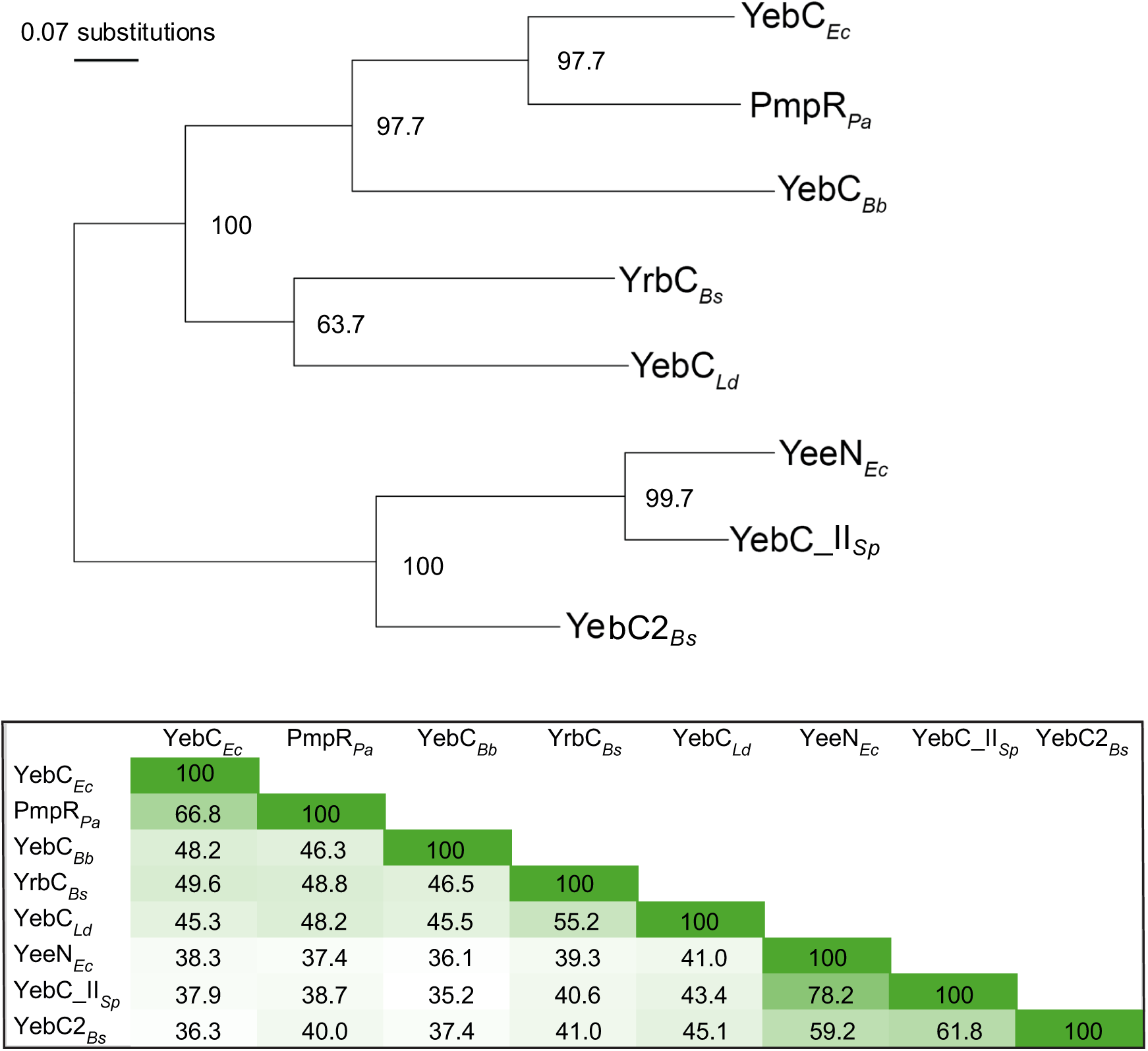
Midpoint rooted maximum-likelihood tree (top) and sequence similarity matrix of characterized YebC family proteins (bottom). Characterized YebC family proteins are labelled with their given gene name and respective organism: *Bs, Bacillus subtilis; Ld Lactobacillus delbrueckii; Pa, Pseudomonas aeruginosa; Ec, Escherichia coli; Bb, Borrelia burgdorferi; Sp, Streptococcus pyogenes.* (Top) Bootstrap values are listed at each node. (Bottom) Pairwise percent identities for the proteins are listed and shaded relative to their homology.

